# MitoTNT: Mitochondrial Temporal Network Tracking for 4D live-cell fluorescence microscopy data

**DOI:** 10.1101/2022.08.16.504049

**Authors:** Zichen Wang, Parth Natekar, Challana Tea, Sharon Tamir, Hiroyuki Hakozaki, Johannes Schöneberg

## Abstract

Mitochondria form a network in the cell that rapidly changes through fission, fusion, and motility. This four-dimensional (4D, x,y,z,time) temporal network has only recently been made accessible through advanced imaging methods such as lattice light-sheet microscopy. Quantitative analysis tools for the resulting datasets however have been lacking. Here we present MitoTNT, the first-in-class software for <u>Mito</u>chondrial <u>T</u>emporal <u>N</u>etwork <u>T</u>racking in 4D live-cell fluorescence microscopy data. MitoTNT uses spatial proximity and network topology to compute an optimal tracking. Tracking is >90% accurate in dynamic spatial mitochondria simulations and are in agreement with published motility results in vitro. Using MitoTNT, we reveal correlated mitochondrial movement patterns, local fission and fusion fingerprints, asymmetric fission and fusion dynamics, cross-network transport patterns, and network-level responses to pharmacological manipulations. MitoTNT is implemented in python with a JupyterLab interface. The extendable and user-friendly design aims at making temporal network tracking accessible to the wider mitochondria community.

## Introduction

Mitochondria are membranous organelles in cells that provide up to 90% of the cellular energy, and are thus fundamental to almost all processes of life from inheriting genetic information to retaining molecular order^1,2^. In mitochondrial diseases, the function of mitochondria is impacted, leading to diminished energy production and cell and organ dysfunction. This is particularly true in high-energy demand organs such as the muscles, heart, and brain. A vast array of diseases such as metabolic disorders^3^, developmental disabilities^4^, epilepsy^5^, neurodegenerative disease^6–8^, cancer^2,9^, and aging^10,11^ may result from mitochondrial dysfunction. Progress in developing pharmacological modulation of mitochondria has been limited, potentially due to the current difficulty in quantitatively measuring the behavior of the cellular mitochondrial network with sufficient spatial and temporal detail.

Measuring the dynamic mitochondrial network is difficult. Far from the solitary kidney bean shapes depicted in many textbooks, interconnected somatic mitochondrial tubules fill all three spatial dimensions and undergo continuous changes in the fourth dimension of time through active and passive motion, fission, and fusion^2^. Conventional fluorescence microscopy technology has been inadequate to simultaneously capture the full spectrum of both mitochondrial morphology and dynamics in all four dimensions (4D). The advent of high-framerate low-phototoxicity fluorescence microscopes such as lattice light-sheet microscopy^12,13^ (LLSM) has now made the detailed 4D characterization of temporal mitochondrial networks possible. Quantitative analysis of this data remains a problem however.

The majority of existing quantitative analysis software was designed for two-dimensional (2D) fluorescence images of mitochondria (MyToe^14^, MitoSPT^15^, QuoVadoPro^16^). For three-dimensional (3D) fluorescence images, MitoGraph^17–19^ is a unique tool for the segmentation and quantitation of 3D mitochondrial network morphology, yet lacks temporal analysis. The software packages TrackMate^20^ and Mitometer^21^ can operate on 4D time-lapse fluorescence microscopy data by performing center-of-mass tracking. However, the abstraction of every mitochondrial fragment as single object poses limitations for accurate sub-fragment level information and network tracking.

Here we present MitoTNT, the first-in-class software for the tracking of the 4D mitochondrial network. MitoTNT builds on the established tools MitoGraph^17–19^ for segmentation and ChimeraX^22,23^ for intuitive visualization. Mitochondria tracking is achieved by solving a linear assignment problem (LAP) that utilizes both spatial and network topology information. Tracking accuracy was validated both in-silico and in-vitro. A reaction-diffusion simulation of the mitochondrial network was created to provide in-silico ground truth for testing. In vitro data of mitochondrial networks was created using LLSM in human induced pluripotent stem cells (hiPSCs). We demonstrate that MitoTNT’s high-resolution mitochondria network tracking is accurate and provides an unprecedented level of detail for mitochondria motility measurement, fission/fusion event detection, and temporal network analysis.

## Results

### Preserved topology enables 4D mitochondrial network tracking

Our first aim was to confirm that high-framerate fluorescence imaging of the 4D mitochondrial network retains enough information for reliable tracking. The somatic mitochondrial networks of tall cuboid hiPSCs were used as a model system (Fig. 1a). LLSM was used to acquire imaging volumes at 3.2s per volume. After deskewing and deconvolution, individual cells were computationally segmented based on the plasma membrane signal (Fig. 1b and Supplementary Fig. 1). MitoGraph^17–19^ was then used to segment the mitochondrial network for consecutive imaging volumes (Fig. 1c,d). At 3.2s frame interval, we observed that changes of the 4D mitochondrial network are predominantly limited to small movements and remodeling events while the overall network structure appeared to be conserved from frame to frame (Fig. 1e). We then quantified this conservation at several acquisition frame rates by applying the scale-invariant feature transform (SIFT)^24^. For small time intervals, SIFT was able to correctly assign network features between frames (Fig. 1f top), but failed for longer time intervals (Fig. 1f bottom). We found that at high volumetric frame rates, mitochondrial network topology is preserved (Fig. 1g). In the next section, we aim to use this conserved temporal information to achieve 4D mitochondrial network tracking.

**Figure 1.**
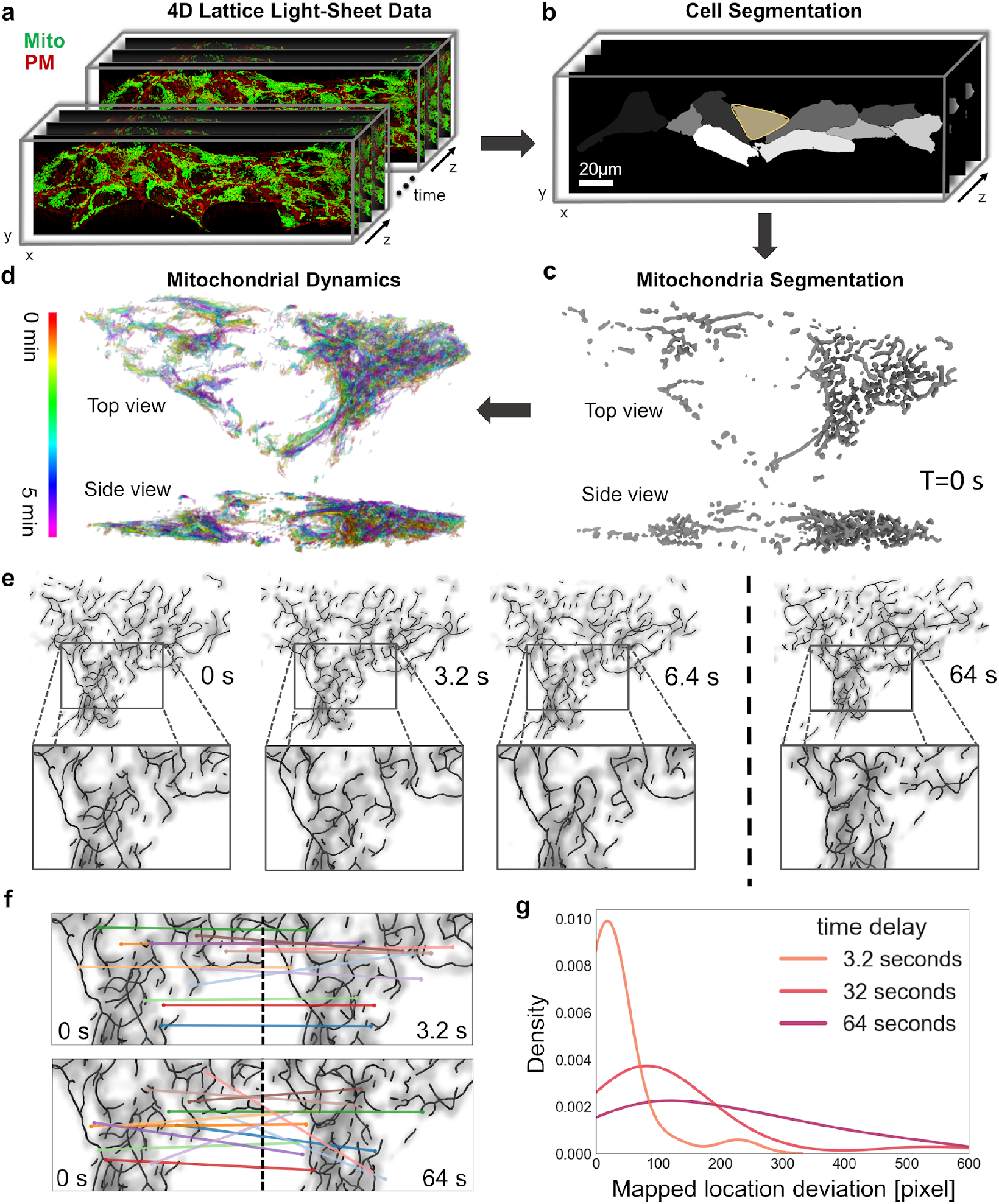
Mitochondrial network topology is preserved in high-framerate 4D fluorescence microscopy data. **a**, Representative 4D (3D+time) lattice light-sheet microscopy data of a hiPSC colony labeled with MitoTracker (mitochondria, green) and expressing CAAX-RFP (plasma membrane, red). **b**, Individual cells in the colony are segmented based on the plasma membrane marker. **c**, Mitochondria fluorescence signal in a single cell is segmented using MitoGraph^17–19^. **d**, Mitochondrial network skeleton dynamics over 5 min every 6.4s (time red to purple). **e**, Mitochondrial fluorescence density and segmented network skeleton are overlaid and shown for frame numbers 0,1,2,20 at frame interval 3.2s. **f**, Scale-invariant feature transform (SIFT) maps image features for two frames separated by 3.2s (top), and 64s (bottom). **g**, Pixel deviation between SIFT-mapped feature locations at different time intervals.

### 4D mitochondrial network tracking using spatial and topological optimization

We formulated 4D mitochondrial temporal network tracking as an optimization problem that uses information preserved in consecutive frames. We chose equally spaced nodes along the mitochondria skeleton as the fundamental units that are tracked (Fig. 2a). Such a discretization of the mitochondrial network can be automatically calculated using the software MitoGraph^17–19^.

**Figure 2.**
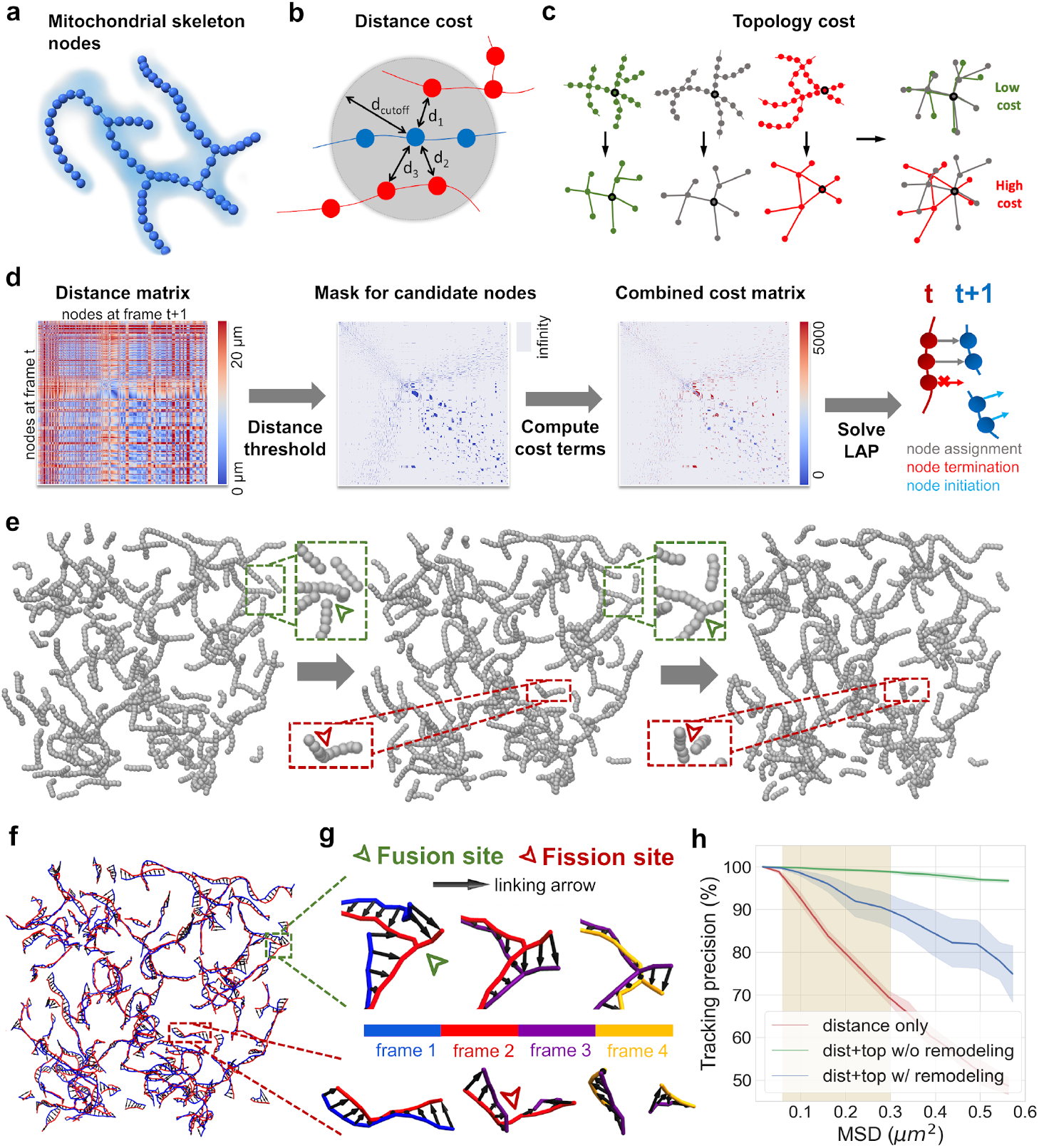
Algorithm design and in-silico validation of 4D mitochondria network tracking. **a**, Discretized nodes along the segmented mitochondria skeleton serve as the basis for network tracking. Cloud: fluorescence density; cylinder: segmented skeleton; sphere: skeleton node. **b-c**, Cost terms used for the linear assignment problem (LAP) formulation of node tracking. Spatial proximity is measured as distances between nodes within two consecutive frames. Topology cost is computed using a graph comparison that assigns low cost for similar local topology. **d**, LAP formulation of node tracking for the mitochondrial network. From left to right: 1) pairwise distance matrix for nodes at frames T and T+1; 2) thresholds to eliminate nodes too far to be tracked; 3) spatial separation and network topology constraints; 4) the solution to the LAP yields the tracking results as linked node pairs, along with terminated and initiated nodes. **e**, Three consecutive frames of a reaction-diffusion mitochondrial network simulation with representative fusion (green) and fission (red) events. **f**, Temporal network tracking for the simulated mitochondria for two consecutive timepoints (blue skeleton: frame 1, red skeleton: frame 2, black arrows: node tracking). **g**, magnification of example in-silico fusion (green triangle) and fission (red triangle) events in **f)**. **h**, Accuracy of in-silico tracking compared to node mean squared displacement (MSD), N=10 simulations. MSD relates to frame as *MSD = 6Dτ*. Commonly achievable frame rates with LLSM highlighted in yellow.

In our previous observation, we found that spatial proximity (Fig. 2b) and network topology (Fig. 2c) are conserved characteristics that likely allow temporal tracking. At high framerates, mitochondrial motion is limited, and the nodes located close to the current position in the next frame tend to be the correct candidates. However, this distance metric quickly decorrelates in dense network regions. Similarly, the mitochondrial network topology remains relatively stable at high framerates and only decorrelates at high fission/fusion rates of the network. We developed a topological dissimilarity score to capture this parameter. The score is computed using a fast alignment-based graph comparison method (see Supplementary Note 4) to measure how different the network topologies around any two candidate nodes are. Nodes embedded in a similar local network topology are more likely to be linked in time.

Similar to established particle/object tracking methods^20,21,25^, we formulated the network tracking problem as a linear assignment problem (LAP) that solves for the optimal node assignment through constraints (Fig. 2d). First, the distances between nodes in two consecutive frames T, T+1 were computed as a pairwise distance matrix. Next, local distance thresholds were estimated for each node at frame T (see Supplementary Note 3). Nodes located within these thresholds at frame T+1 were considered candidate nodes while those beyond were ignored. Then, network topology was incorporated using the topological dissimilarity score for each candidate node pair (node at T and candidate node at T+1). The distance and topology costs were then combined with equal weights. Mitochondrial dynamics and imaging artifacts often contribute to fluctuations in the number of skeleton nodes. To account for this fluctuation, we added additional constraints to the final cost matrix (see Supplementary Note 3), thereby permitting three options for a temporal assignment: 1) link two nodes between frames, 2) terminate a node in the current frame, or 3) initiate a new node in the next frame. Finally, the frame-to-frame tracking result is given as the optimal node assignment to the LAP by minimizing the global sum of the final cost matrix. Gap closing is performed at the end of frame-to-frame tracking in order to connect prematurely terminated node tracks, using the same cost terms (Supplementary Note 5).

### In-silico validation of MitoTNT through spatial reaction-diffusion simulations of mitochondrial networks

Our next aim was to validate our tracking algorithm using synthetic data as ground-truth. A meso-scale reaction-diffusion simulation was developed to model temporal mitochondrial networks (Fig. 2e). We used the ReaDDy^26,27^ framework to model mitochondria as connected mitochondrial skeleton particles that were held together by bond, angle, and repulsion potentials. Mitochondrial motion was assumed to be diffusive only. The spatial distribution and density of the in-silico mitochondrial network was modeled after in-vitro imaged mitochondrial networks that we found to resemble a mixture of Erdös–Rényi random networks (Supplementary Fig. 4a-b). Fission and fusion were included as structural reactions such that fission reactions remove a bond between skeleton nodes and fusion reactions create a bond between unbound skeleton nodes. Experimental observations of fission and fusion rates were adjusted through iterative sampling of fission and fusion reaction rates (see Supplementary Note 5).

Tracking accuracy of our algorithm was subsequently tested using this simulation as ground-truth. We found that each fragment of the mitochondrial network, as well as fission and fusion events are tracked faithfully with few mis-assignments (See Fig. 2f,g). We found that the distance constraint alone results in relatively poor tracking performance (Fig. 2h, red curve) likely due to ambiguous assignments in the dense mitochondrial network. In contrast, when paired with the topology constraint, consistently reliable tracking was achieved with fission and fusion switched off (95-100% accuracy, Fig. 2h green) or on (> 90% accuracy, Fig. 2h blue) in the regime relevant for LLSM (shaded region, see Supplementary Note 6).

### In-vitro validation and evaluation of 4D mitochondrial network tracking

We next validated our tracking algorithm on LLSM data of 4D mitochondrial networks in cultured cells. CAAX-RFP hiPSC colonies were labeled with MitoTracker Green and imaged at 3.2 seconds per volumetric frame for a duration of 5 minutes. Cells and mitochondrial network were segmented (Fig. 3a-b) and MitoTNT used to track the network (Fig. 3c and Movie 1). Careful examination of the tracking results showed that the mitochondrial network skeleton is faithfully tracked over time (see Fig. 3d). Depending on the level of granularity required for the biological question of interest, tracks for the nodes (Fig. 3e) that belong to the same segment (Fig. 3f), or the same fragment (Fig. 3g) can be obtained. We found that somatic mitochondrial motility is diffusive not only on the fragment-level but also on the mitochondrial skeleton node-level (Fig. 3h, 3i, S6). We observed, that the high-resolution tracking on the level of mitochondrial skeleton nodes illustrates that mitochondrial motility and dynamics exhibit complex spatial and temporal details and a heterogeneity in speed and orientation (Fig 3e-g, lower panel). Finally, we compared our high-resolution tracking results to previously published values of lower-resolution center-of-mass tracking. We found that our average mitochondrial network fragment motility for hiPSC mitochondria (0.06±0.03 μm/s) is in good agreement with motility data from 3D spheroids (0.03μm/s) and 2D adherent cells (0.08μm/s) (Supplementary Figure 6).

**Figure 3.**
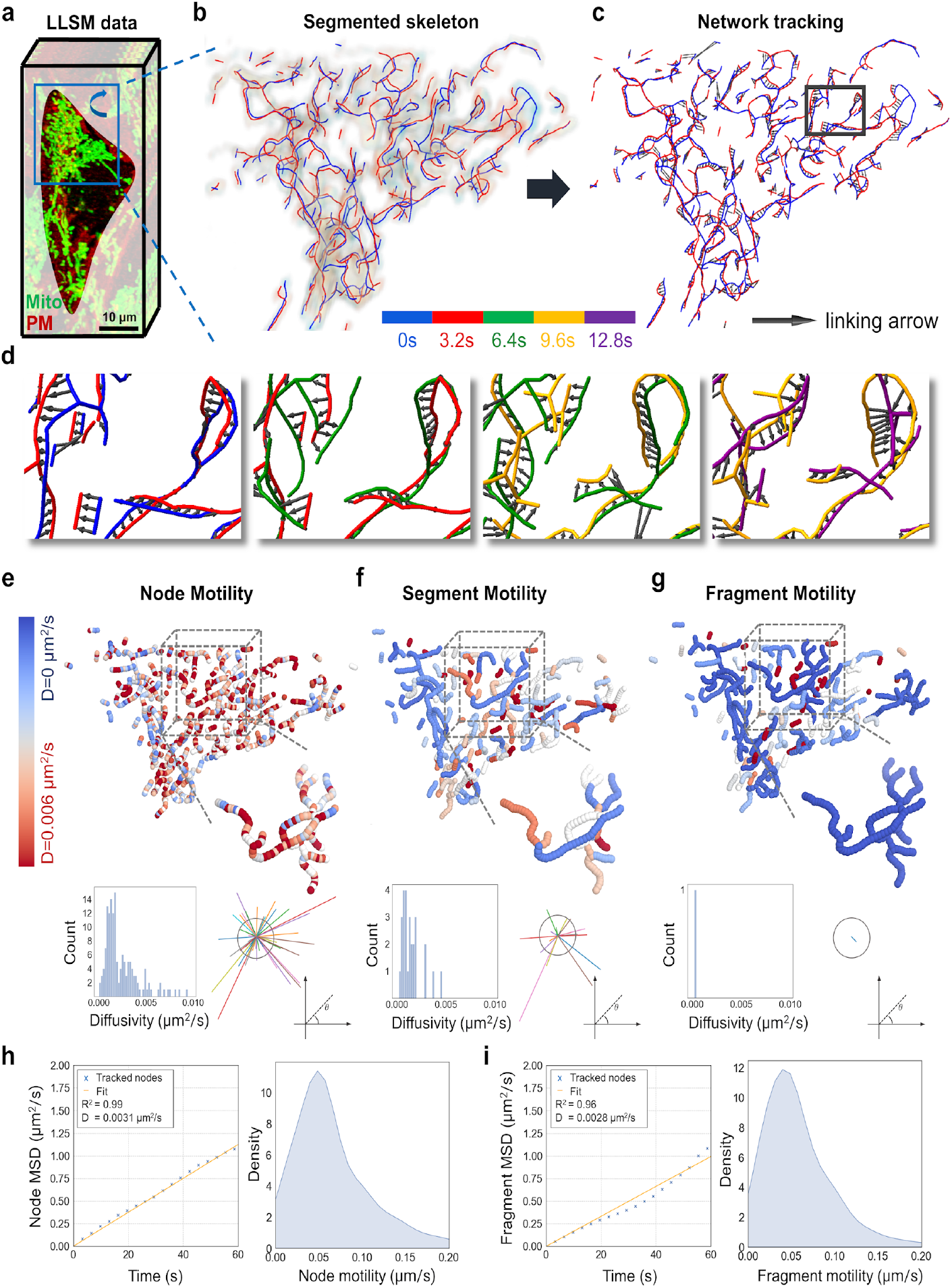
In-vitro validation and evaluation of 4D mitochondria network tracking. **a**, LLSM volumetric snapshot of a segmented cell. Green: mitochondrial network. Red: plasma membrane. **b**, Zoom-in on the mitochondrial network in **a)**. Fluorescence signal and segmented network skeleton are overlaid for two consecutive frames (blue: 0s, red: 3.2s). **c**, Tracking of the network nodes for the two frames in **b)** visualized by black arrows. **d**, Zoom-in to a representative region (box) in **c)** tracked over 12.8s. The skeletons are colored in blue, red, green, yellow, and purple in the order of time. See also Movie 1. **e-g**, Top: Mitochondrial nodes are colored by diffusivity at node, segment, or fragment levels from high (red) to low (blue) diffusivity. Bottom left: Distribution of diffusivity values, bottom right: linking vectors compared to a fixed reference vector. **h-i**, MSD curve and motility distribution for nodes **h)** and fragments **i)**.

### High-resolution mitochondria tracking reveals heterogeneous sub-fragment motility and correlated movement patterns

We observed individual fragments displaying a wide range of movement patterns that include translational, and rotational components. Branches of the same mitochondrial fragment can simultaneously undergo motions with different orientations and modes. Here we showcased three examples: 1) a small fragment exhibiting twisting motion (Fig. 4a), 2) a medium-sized fragment exhibiting concentric inward motion (Fig. 4b), and 3) a large fragment exhibiting convolution of different motility patterns (Fig. 4c).

**Figure 4.**
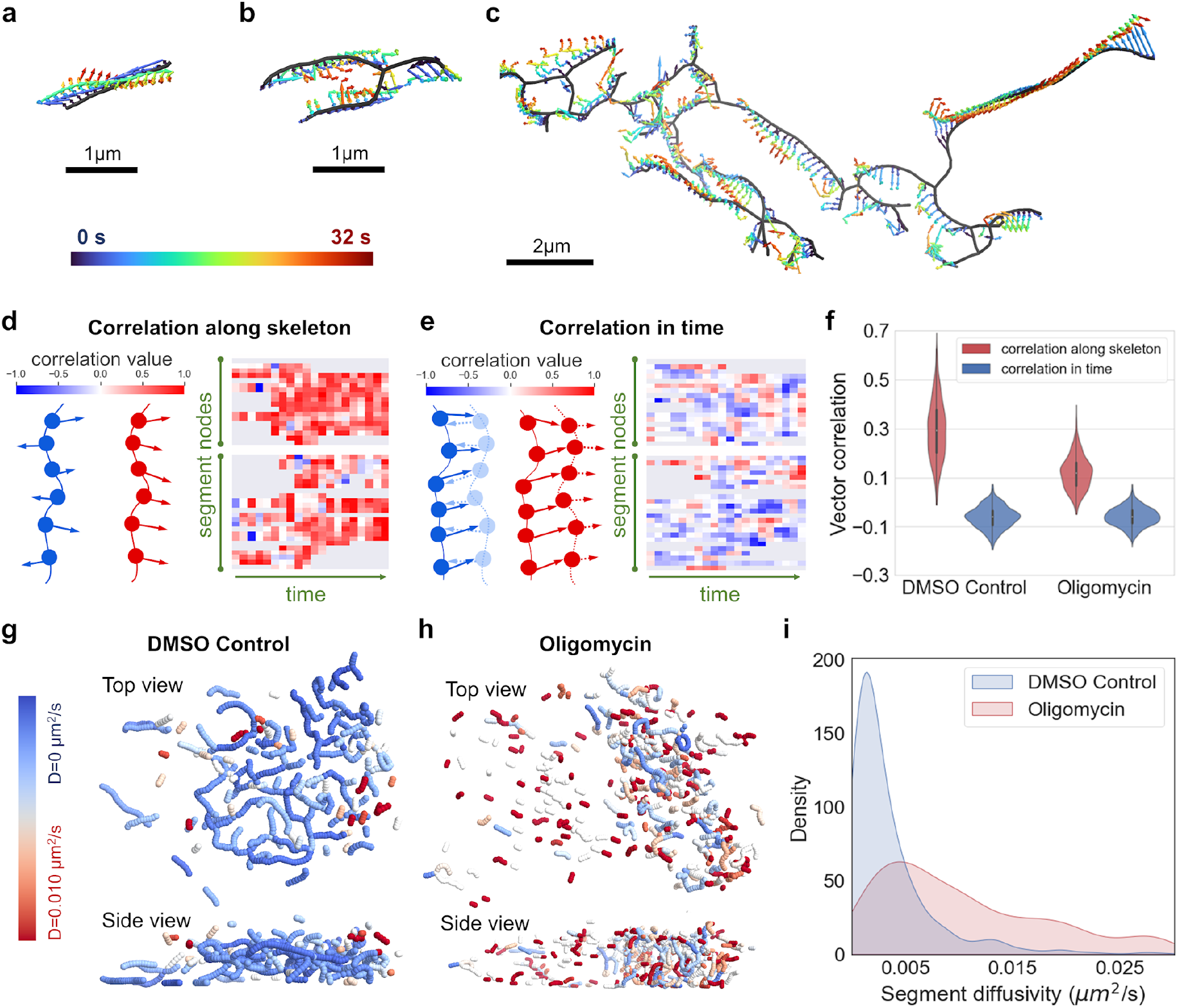
Mitochondrial network motility analysis. **a-c,** Tracking of three representative mitochondrial network fragments for 32 seconds (time blue to red). **a**, A small fragment displays twisting motion. **b**, A medium-size fragment displays inward motion. **c**, A large fragment displays complex motion patterns. **d**, spatial tracking vector correlations between neighboring nodes. Left: illustration. Right: Heatmap of vector correlation for segment nodes (columns) at different timepoints (rows). **e**, temporal tracking vector correlation for the same node at consecutive frames. Left: illustration. Right: Heatmap of correlation values for segment nodes (columns) at different timepoints (rows). **f**, Violin plot of spatial (red) and temporal (blue) correlation values between control and oligomycin-treated cells. **g-h**, Spatial structure of somatic mitochondrial network overlayed with mitochondria segment diffusivity in control and oligomycin-treated cells. **i**, Kernel-smoothed distribution of segment diffusivity for 2552 segments in control cells (blue), and 2376 segments in oligomycin-treated cells (red).

To further investigate network branch motility, we correlated tracking vectors between adjacent nodes on the same segment, and between the same node at consecutive frames. We observed that spatial correlation along the segment skeleton is predominantly positive (Fig. 4d) demonstrating a concerted motion. In contrast, temporal correlation between frames is predominantly zero (random motion) to slightly negative (oscillating motion), while interspersed with short period of positive values (directional motion) (Fig. 4e). This data confirms that mitochondrial branches move as a unit, but in a relatively random manner (Fig. 4f, control).

We employed the ATP synthase inhibitor oligomycin that induces mitochondrial fragmentation^28^ to investigate if our motion correlation findings are dependent upon network morphologies (Fig. 4g,h). We observed that while temporal motion correlation remains similar, the spatial motion correlation dropped. Furthermore, we observed that drug induced fragments move considerably faster compared to control (Fig. 4i).

### Mitochondrial network tracking reveals local fission and fusion fingerprints and asymmetric fission and fusion preferences

Our high-resolution network tracking allows us to precisely locate fission and fusion events in the mitochondrial network with sub-fragment spatial resolution and high temporal fidelity (Figs. 5a S5). To provide mechanistic insights into network remodeling, we compared the motility between randomly selected nodes and nodes undergoing fission and fusion. We observed that diffusivity for nodes undergoing fission and fusion is nearly two times the diffusivity for randomly chosen nodes (Fig. 5b). This data suggests that mitochondrial fission and fusion remodeling might involve local rearrangements at the event site as suggested previously^29^.

**Figure 5.**
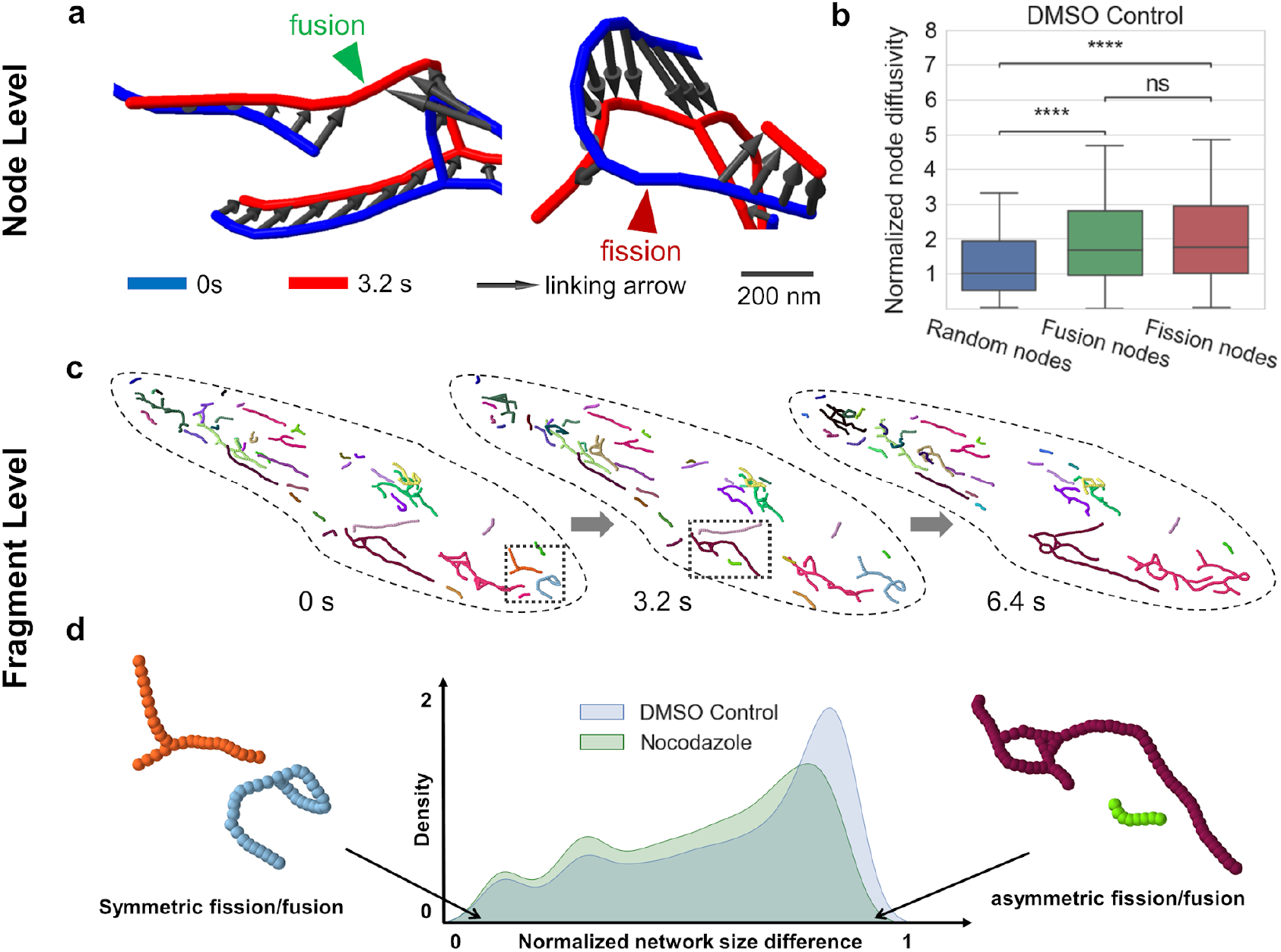
Mitochondrial network remodeling analysis. **a**, Representative snapshots of tracked fusion event (left), and fission event (right). **b**, Node diffusivity is significantly lower for randomly selected nodes (blue) as compared to nodes undergoing fusion (green) and fission (red). Student’s t-test used, and p-values are 6.565E-25, and 1.237E-24, for random vs. fusion nodes and random vs. fission nodes, respectively. **c**, Representative tracking of mitochondrial fragments in hiPSCs over three timepoints (one color per fragment) **d**, Analysis of fission/fusion preferences with respect to fragment size shows that asymmetric fission/fusion events are more likely to occur. This pattern is less pronounced but preserved in nocodazole-treated hiPSCs with disrupted microtubules.

Based on node tracking, each individual mitochondrial network fragment can be tracked (Fig. 5c) and the selectivity of fission and fusion events recorded in terms of fragment size. For each fission or fusion event, the normalized network fragment size difference was computed, with values close to 0 corresponding to a symmetric fission/fusion (Fig. 5d, left), and values close to 1 indicating fragments of drastically different sizes (asymmetric fission/fusion) (Fig. 5d, right). We found that there is a significant portion of asymmetric fission/fusion events (Fig. 5d, blue). Asymmetric fission/fusion events between large healthy mitochondria and small unhealthy mitochondria have been proposed to separate dysfunctional mitochondria targeted for mitophagy, or rescue damaged mitochondria by supplying essential materials^30,31^. We hypothesized that this dynamic selectivity bias is facilitated by the cytoskeleton. In cells treated with 10 μM of nocodazole to disrupt microtubules, we observed a decrease in asymmetric fission/fusion (Fig. 5d, green). This observation points to a potential role of cytoskeleton in mediating selective fission/fusion as has previously been suggested^32^.

### 4D mitochondrial network tracking shows drug-dependent network remodeling rates, network transport, and network resiliency

4D mitochondrial network tracking allowed us to investigate the mitochondrial network from the perspective of a graph temporal network (Fig. 6a). Specifically, it is now possible to quantify a) remodeling of the mitochondrial network, b) flux across the network as it moves spatially and is being remodeled, and c) resiliency of the network to damage.

**Figure 6.**
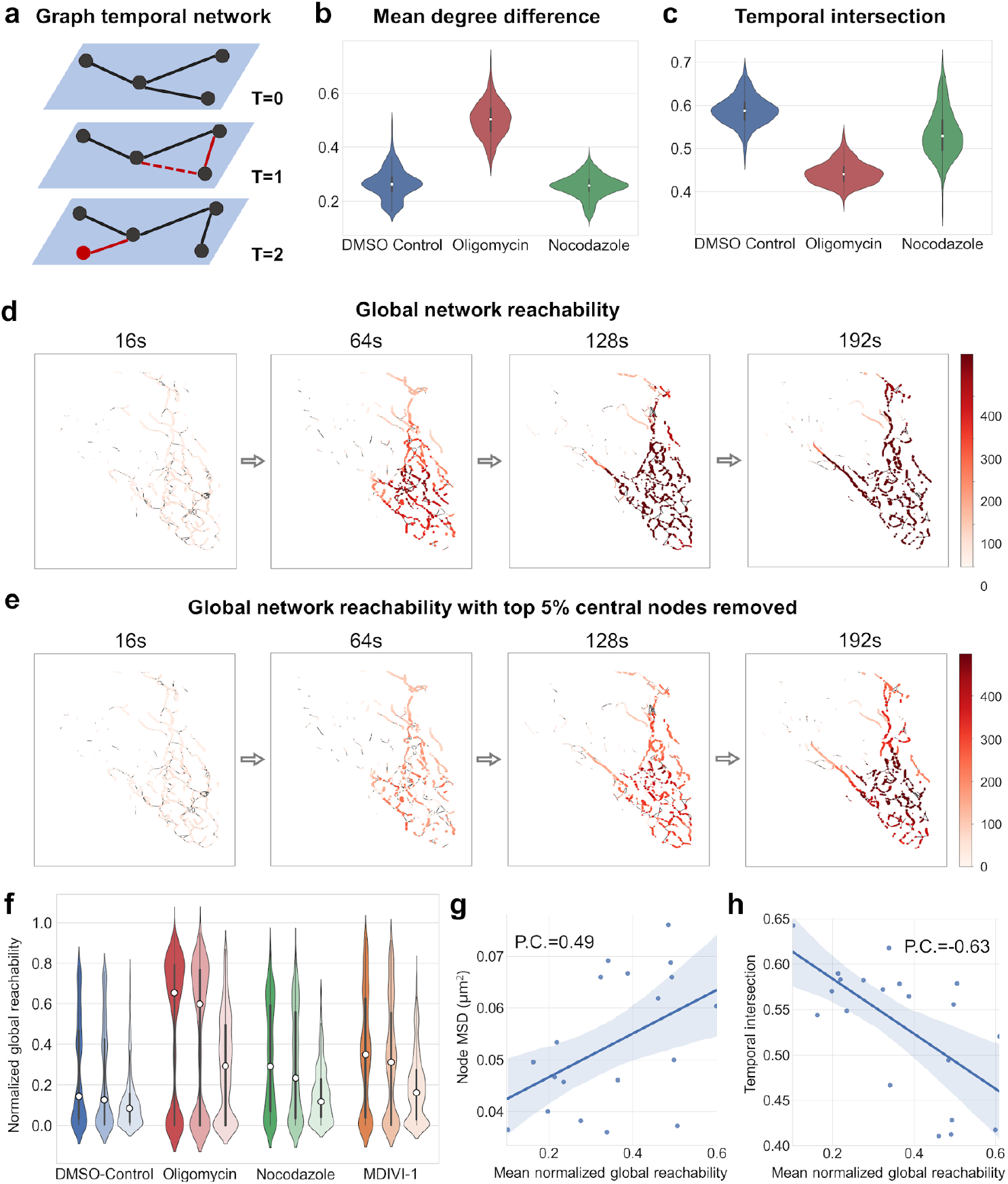
Temporal characteristics of mitochondrial network remodeling, flux, and damage resilience. **a**, Temporal networks display node and edge dynamics that have an influence on network transport and resilience (newly added or removed nodes/edges highlighted in red). **b**, Mean degree difference between control, oligomycin, and nocodazole. **c**, Temporal intersection between control, oligomycin, and nocodazole. **d**, Global network reachability in a representative somatic mitochondrial network depicted as a color gradient (dark: high reachability, light: low reachability). **e**, Global network reachability where the top 5% highest betweenness-centrality nodes were removed. **f**, Mean normalized global reachability in different drug induced conditions. Triplets indicate no nodes removed (left), 5% random nodes removed (middle), and 5% most connected nodes removed (right). **g**,**h**, Correlation of network reachability with node MSD and temporal intersection.

To quantify mitochondrial network remodeling, the mean degree difference (DD) and the temporal intersection (TI) were calculated. A low DD indicates a low rate of nodes breaking off from their neighbors and reconnecting with other nodes. Inversely, A low TI implies that the network is very dynamic and does undergo drastic remodeling. We found that control hiPSCs showed a low DD of 0.28 (Fig. 6b) and a high TI 0.58 (Fig. 6c), indicating that the network is relatively stable with relatively little turnover. In contrast, when treated with oligomycin, we observed a 0.52 DD and 0.44 TI indicating a high level of network remodeling. We hypothesized that cytoskeleton influences drive network remodeling events. However, treatment with nocodazole did not induce drastic changes in neither metric for network remodeling (Fig. 6b,c).

To quantify transport across the 4D mitochondrial network, we simulated a random walk on the tracked temporal mitochondrial networks and measured the process in the form of network reachability (see Supplementary Note 9). Reachability for a node indicates how easily can material/information reach this node from various parts of the overall network, via the time-respecting paths defined by the network tracking. In control conditions, we observed that almost every part of the network was in reach within ~120s (Fig. 6d) and that network nodes showed a low average reachability of 0.18 (Fig. 6f). Comparatively higher reachability was reached with nocodazole (0.29), mdivi-1 (0.35), and in particular with oligomycin (0.64).

To quantify mitochondrial network resiliency, mitochondrial node reachability was calculated in networks where the top 5% of highest connected nodes were removed, as measured by betweenness of centrality. We observed that the global reachability for a large number of nodes was significantly reduced, particularly those isolated from the larger well-connected fragments. This observation suggests certain central nodes may be essential to the material and information transport within the cellular mitochondrial network (Fig. 6f). To quantify the relationship between network motility, remodeling, and reachability, we calculated the Pearson’s correlation coefficients between the mean normalized global reachability, the mean node displacement (Fig. 6g), and the node TI (Fig. 6h). The positive correlation with node displacement, and negative correlation with TI suggests that long-range movements and enhanced network remodeling both lead to quicker percolation through the network.

## Discussion

Here we presented MitoTNT, the first-in-class software for mitochondrial temporal network tracking in 4D volumetric fluorescence microscopy data. Recent advances in low phototoxicity volumetric live cell imaging allow fast high-resolution acquisition of the somatic mitochondrial temporal network. Now, MitoTNT allows the automated tracking of this temporal network for the first time. Based on mitochondria skeleton segmentation and discretization through MitoGraph, MitoTNT solves the linking problem of discretized mitochondria skeleton nodes through time. An efficient, alignment-based graph comparison algorithm was used to capture network topology information and pair it with distance constraints for temporal linking. Tracking was validated using both in-silico and in-vitro methods. We created polymer-based spatial mitochondrial simulations that include fission and fusion reactions and are parameterized to reproduce experimental observations to quantify the tracking fidelity of our algorithm. We found that MitoTNT performs with >90% tracking accuracy on these datasets. When comparing tracking performance on experimental in-vitro datasets, we found that MitoTNT faithfully tracks the 4D mitochondrial network and reproduces experimental observables such as mitochondrial diffusivity and speed as compared to published values in the literature. Based on fluorescence microscopy and computational image segmentation, MitoTNT is limited by a microscope’s ability to record high signal to noise volumetric images of the mitochondrial network to ensure high quality segmentation. Future efforts might use advances in machine learning^33–35^ to improve segmentation quality and reliability.

We highlighted three applications of MitoTNT: 1) high-resolution mitochondria network motility analysis, 2) node-level mitochondrial fission/fusion analysis, and 3) mitochondria temporal network analysis. For motility analysis, we showed that the previously hidden complexity of sub-fragment motility can now be characterized. By coupling network sub-compartment motility with other mitochondrial fluorescence readouts (e.g., membrane potential, reactive oxygen species, mtDNA nucleoid), future studies employing network tracking will have the potential to investigate the functional aspects of mitochondrial motion in cellular physiology.

For node-level fission/fusion analysis, we showed that mitochondrial fission and fusion dynamics can be registered at sub-fragment resolution. Compared to fission/fusion detection for object-based tracking, our approach is highly versatile in distinguishing sub-types of mitochondrial remodeling events such as kiss-and-run events, sustained fission/fusion events, intra-fragment events, and inter-fragment events. We predict, that the high spatio-temporal resolution offered through mitochondrial network tracking will become instrumental in studying selective fission/fusion^32^ and mitochondrial quality control^7,36^.

The characterization of somatic mitochondrial networks as temporal networks through MitoTNT now allows using the full power of mathematical models for graph temporal networks for mitochondria analysis, for example determining community formation within the mitochondrial network, understanding the efficiency of metabolic flow, or characterizing various cell types and states using network motifs. By combining 4D fluorescence imaging, network tracking, and functional simulation, cellular metabolic state profiling based on microscopy data can now be conducted, opening the door for high-content screening of such states. We hope that MitoTNT’s extendable software design and open-source code availability will contribute to forming a community around mitochondria temporal network tracking and will allow the field to quickly explore the indicated directions.

## Methods

### Human induced pluripotent stem cell (hiPSC) culture

All studies involving hiPSCs were performed under approval from the University of California San Diego IRB/ESCRO committee. WTC hiPSCs expressing the CAAX domain of K-ras tagged with mTagRFP-T were created at the Allen Institute for Cell Science and obtained through the UCB Cell Culture Facility. hiPSC colonies were expanded on dishes coated with growth factor-reduced Matrigel (Corning, 354230) in mTeSR1 (Stemcell Technologies, 85850) containing 1% penicillin/streptomycin (Gibco, 15140122). Colonies were washed with DPBS (Gibco, 14190144) and detached with accutase (Stemcell Technologies, 07920) before plating onto imaging dishes. Cultures were tested routinely for mycoplasma.

### Drug treatments

All drugs were dissolved in DMSO to make a stock solution and diluted in PBS to prepare a 100X working stock. Cells were treated with oligomycin (20uM, 2 hr), nocodazole (5 uM, 30 min), and MDIVI-1 (10 uM, 12 hr) without wash.

### Live cell imaging

CAAX-RFP hiPSCs were stained with 100 nM MitoTracker Green FM (Invitrogen, M7514) for 30 min prior to imaging. Cells were plated onto 25 mm MatTek dishes and imaged in phenol-red free mTeSR1 (Stemcell Technologies, 85850). Cells were kept under 5% CO_2_ and 37 degrees C. For imaging, we used Zeiss LLSM 7 with 10× N.A. 0.4 illumination objective lens and 48× N.A. 1.0 detection objective lens. We acquired images in two channels: green channel with excitation at 488nm and emission at 512nm; red channel with excitation at 561nm and emission at 597nm. For both channels we used 18% laser power and 8ms exposure. The illumination light-sheet was the Sinc3 beam with length 15 μm, thickness 650 nm and no side lobes. The volume size was 2048 × 448 × 57 pixels or 296.94 × 64.96 × 8.12 μm with isotropic pixel size 145 nm after coverglass transformation. Images were saved with bit depth 16 bits. For each region, we imaged 93 frames with frame rate 3.26 s per volume for total 5 min. For LLSM data processing, we used the Lattice Lightsheet 7 Processing Module on ZEN Blue for deconvolution, deskew, and cover glass transformation. Further processing is then done using MitoGraph and MitoTNT.

## Code availability

Please find links to documentation, source code and other information at https://www.mitotnt.org. Specifically, code, installation guide, and sample data are available at https://github.com/pylattice/mitoTNT.

## Acknowledgements

The authors thank S. Rafelski, G. Pekkurnaz, and E. Koslover for helpful discussions and early feedback on the manuscript. This study was supported by funds from the Hartwell Foundation through an Individual Biomedical Research Award to J.S.

## Contributions

Z.W., P.N., and J.S. conceived of the project; Z.W. developed the tracking tools, performed the modeling, cell culture, and imaging; P.N. developed and implemented the graph-based tools; C.T. developed the cell lines and performed cell culture and imaging; S.T. performed cell culture; H.H. performed the imaging, Z.W., P.N., and J.S. wrote the manuscript.

