## Supplementary Information for "MitoTNT: Mitochondrial Temporal Network Tracking for 4D live-cell fluorescence microscopy data"

### Supplementary materials

Zichen Wang[1,2], Parth Natekar[1,2], Challana Tea[1,2], Sharon Tamir[1,2], Hiroyuki Hakozaki[1,2], Johannes Schöneberg[1,2,*]

[1] Department of Pharmacology, University of California, San Diego, San Diego, CA, 92093
[2] Department of Chemistry and Biochemistry, University of California, San Diego, San Diego, CA, 92093

* Address correspondence to

Johannes Schöneberg, Ph.D.
Departments of Pharmacology and Chemistry and Biochemistry
University of California, San Diego
Center for Neural Circuits and Behavior, Room 106B
La Jolla, CA 92093


**Supplementary Note 1**

**Cell segmentation**

The cell segmentation for individual stem cells in a stem cell colony is performed as follows. The red fluorescence signal from CAAX membrane markers is first normalized and smoothed. We then use the 2D filament filter from Allen Cell & Structure Segmenter[1] to threshold the membrane contour for the middle z-section. By dilating and inverting the membrane contour, seed labels for the watershed segmentation[2] in scikit-image[3] are automatically obtained. In order to ensure that each cell has a single seed label positioned near the center of the cell body, the seed labels are manually corrected in ImageJ[4] if necessary. Next, we use the seed labels from the middle z-section to perform 3D watershed segmentation on the entire z-stack. The resulting cell segmentation masks are again manually checked, and used to crop single cells. Since the cell movement was found to be minimal on our timescales of 5 minutes per movie, we could apply the first frame's segmentation result to the entire movie.

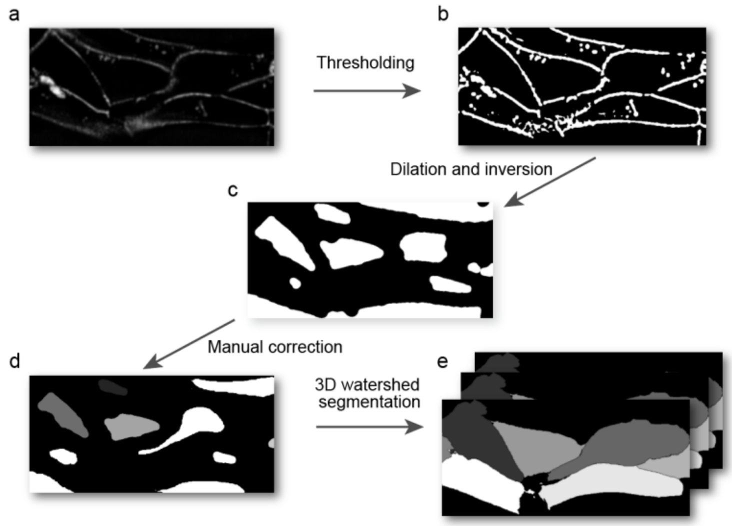


**Supplementary Figure 1. | Cell segmentation workflow.** a, Fluorescence signal from CAAX membrane marker in the middle plane. b, Cell contour is thresholded. c, Center of the cell is highlighted through dilation and color inversion of the cell contour. This is used as the seed for watershed segmentation algorithm. d, The seed is manually checked and corrected. e, The seed for the middle plane is used to segment cell membrane in 3D using the watershed method.

#### Supplementary Note 2

#### Mitochondria segmentation

We used MitoGraph 3.0[5] for mitochondria segmentation. The input for MitoGraph was the fluorescence signal of individual cells segmented in the previous step along with default parameters. Adaptive thresholding with block size of 10 pixels was used to calculate a block-dependent local threshold. The output of MitoGraph includes the segmented mitochondrial network and the segmented mitochondrial skeleton which includes network edges and network node attributes including 3D coordinates, fluorescence intensity, and tubular width. A custom python script was created to read MitoGraph outputs, construct the full-resolution mitochondrial networks, and compile the networks with node attributes as a python-igraph[6] object.

#### Supplementary Note 3

#### Network tracking formulated as a linear assignment problem for node assignment

To track nodes at frames T and T+1, the node coordinates are used to calculate a pairwise distance matrix where element at row m and column n equals the distance between node m at frame T and node n at frame T+1. To limit nodes under consideration for a temporal link, a threshold is implemented. The user can specify the threshold as the distance to the N-th closest neighbor for each node. Additionally, the user can input the maximum node speed, such that nodes located beyond maximum node speed * frame interval are not considered for linking. In our case, we found that the distance to 10$^{th}$ nearest neighbor is sufficient for tracking at relatively high framerate. The graph comparison score (topology cost) is then computed for the candidate node pairs. The combined cost matrix is the product of distance and topology cost matrices, whose weight can be adjusted by the exponents. We used equal weighting with exponents both equal to 1. We then added the self-assignment matrices with alternative cost equal to 98$^{th}$ percentile of all previous assignment cost (2 * the minimum of all potential costs for the first frame). The auxiliary matrix is filled with blocking values. The exact definition of self-assignment matrices and auxiliary matrix can be found in the U-track paper[7]. The final cost matrix is then solved using the Jonker-Volgenant algorithm[8]. The tracks of network nodes are then stored and dynamically updated during frame-to-frame tracking.

**Supplementary Note 4**

**Efficient graph comparison using an alignment-based method**

The topological dissimilarity cost term compares the local network topologies between the nodes from two frames. This is an inexact graph matching problem with partial node-correspondence since only the two nodes for which the cost is to be computed, termed the target nodes, are known. Existing methods including graphlet-based methods [ref], alignment-based methods [ref], spectral methods [ref], and recent portrait divergence methods [ref] are usually designed for large complex networks, computationally non-trivial and application specific. Mitochondrial networks usually have much less convoluted connectivity, and thus a small network neighborhood is usually sufficient for our tracking purposes. The proposed graph comparison algorithm needs to prioritize computational efficiency to accommodate the many iterations over network nodes and timepoints for large LLSM dataset. Thus, we have designed a heuristics-driven, alignment-based method that maps the nodes in two subnetworks, and compute the topological dissimilarity score as the norm of the differences between two mapped adjacency matrices.

We define two types of networks: 1) the classic networks consisting of terminal (degree=1) and branching (degree>2) nodes; 2) the full-resolution networks which include additional bulk nodes (degree=2), equally spaced along the skeleton. We approached network tracking by tracking the all the nodes in the full-resolution networks . We formulate the network tracking problem as a linear assignment problem (LAP) for node assignments under certain spatial and topological constraints.

First, in order to efficiently compare the overall network topology around the target nodes, two selected full-resolution networks are converted to the classic networks by eliminating the bulk nodes and directly connecting the terminal and branching nodes. Second, subgraphs up to a user-defined maximum k-level from the target nodes are used, where the k-th level refers to all the nodes whose shortest path to the target nodes have k nodes. For the subgraphs, breadth-first operations starting from the target nodes are performed to 1) convert the graphs to trees by opening loops, and 2) add pseudo-nodes and pseudo-edges of weight zero to ensure that the numbers of nodes at each level are the same. Third, we define a node dissimilarity score between node pairs, and use this score as the cost in the LAP to solve for the optimal node mapping at a given level. We iterate this step for each level until the maximum level is reached. Now that the graph alignment is complete, the topological dissimilarity score between the target nodes can be given as the Euclidean norm of the differences between the two adjacency matrices with distance weights.

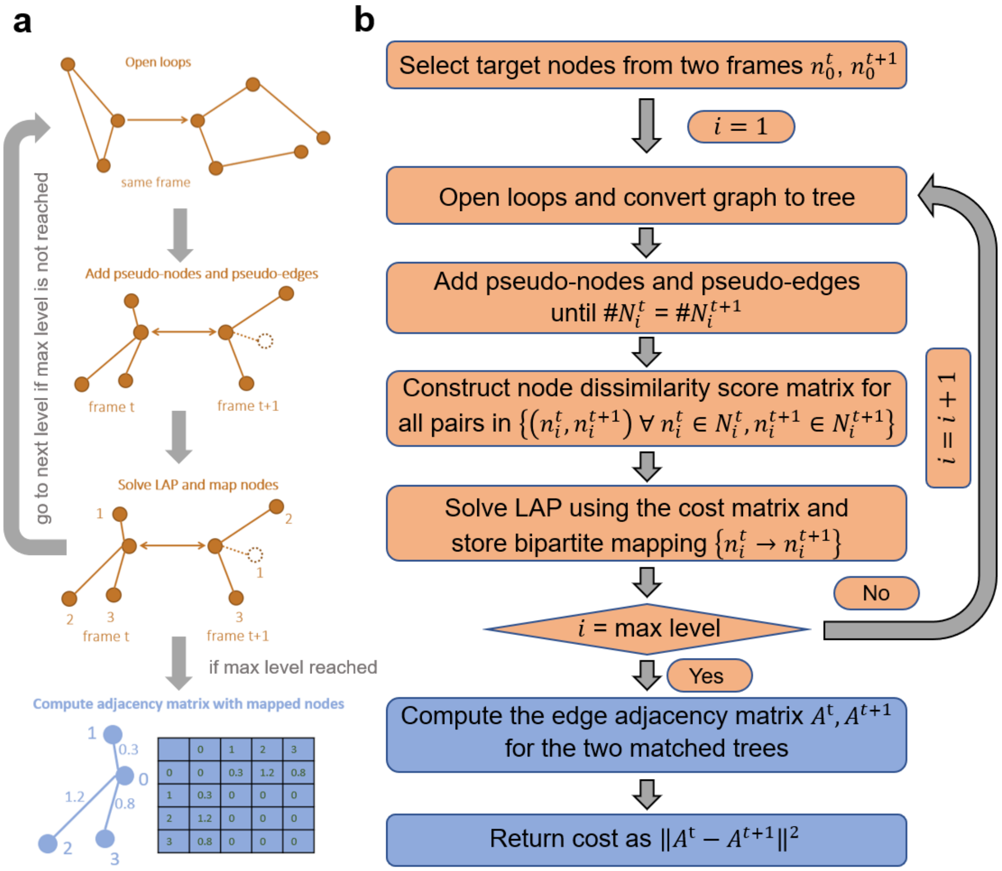


**Supplementary Figure 2. | Illustration for the graph comparison algorithm. a,** Visual illustration for the alignment-based graph comparison algorithm. **b,** Detailed pseudo-code for the algorithm.

**Supplementary Note 5**

**Gap closing**

Once frame-to-frame node tracking is complete, node tracks with frames more than min_track_size are qualified for gap closing. The distance and metric costs are computed between the last frame in track A ($frame_{end}^{A}$) and the first frame in track B ($frame_{start}^{B}$) if both of the following criteria were met: 1<gap size<max_gap_size, where gap size = $frame_{start}^{B}$ - $frame_{end}^{A}$, distance < gap size * 9 * variance(combined track displacements). Because the total number of tracks can be large (more than 10,000), the cost matrix for all tracks can be memory-consuming, and solving the LAP of such a matrix can be computationally expensive. To solve this problem, we divide the full cost matrix into overlapping blocks to obtain sub-optimal global assignment. First, we sort the tracks based on the start frame so that short tracks that can be linked are positioned relatively close. Next, we crop a NxN block at the top left diagonal of the full cost matrix, where N = max_gap_size * average # of tracks per frame, and use this matrix to close gaps for the tracks involved. Finally, we move this block 0.8*N tracks down the diagonal, to allow gap closing for tracks at the boundary. We repeat this process until the full cost matrix was traversed. This iterative gap closing scheme improves memory and computation performance without noticeably changing the number of closed tracks.

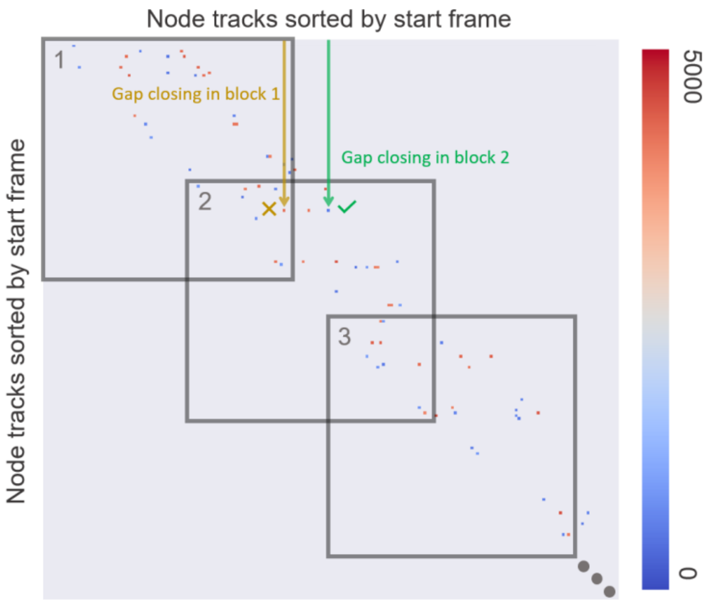


**Supplementary Figure 3. | Memory-efficient gap closing scheme using overlapping cost matrix blocks.** An illustrative gap closing matrix is shown. Row and column indices are node track IDs ranked by the track start frame number. The cost terms are calculated as the product of the distance and topology costs for the end node of the row track, and the start node of the column track. Thus, no assignments are allowed for the lower triangle. Because the number of tracks can be very large, overlapping blocks of the cost matrix are used for track assignments to reduce memory usage and speed up computation. Two track assignments for a track in the overlapped region between two blocks are shown. Because the track is on the edge of the block, the assignment from block 1 has high cost and is sub-optimal (yellow). However, because the block 2 includes more potential tracks, the assignment from block 2 gives the optimal track assignment.

#### Supplementary Note 6

#### Reaction-diffusion simulation of spatial mitochondrial networks

The particle-based reaction-diffusion simulation tool ReaDDy was used to create a synthetic ground truth dataset of 4D mitochondrial dynamics. We setup a rectangular box potential to represent a cell with dimensions of x, y, z = 20, 20, 5 microns with x, y, z = 2000, 2000, 500 units, where each simulation spatial unit equals 10nm. We added a harmonic boundary potential with force constant $k_{box}$=100. Mitochondrial skeleton nodes are represented as individual particles in this box. To form interconnected chains of such particles to represent the mitochondrial network, each node is modeled by the spherical repulsion potential with equilibrium diameter of 350nm, reflecting real-world mitochondrial tubular width[9]. Between each connected node there is a bond potential with $k_{bond}$=1, an angle potential with $k_{angle}$=10, and equilibrium angle=180 degrees. All potentials are harmonic.

To generate the initial classic networks, we combine multiple smaller random networks. For each random network partition, n=60 network nodes are randomly connected with mean degree p=1.0, and any unconnected nodes are removed. More networks are created until the total number of nodes N=300. To convert the classic networks into the full-resolution networks in space, we randomly assign the position for one node in the network, and start traversing the entire network starting from that node. Each edge in the classic networks is replaced by 2-5 bulk nodes. Fission events are modeled by removing a bond potential between connected nodes and fusion reactions are modeled by  event is modeled by topology dissociation reactions. For each connected component, a random edge is deleted at a probability proportional to the total number of edges. The separated components again can undergo fission events, until there are fewer than four nodes to avoid unrealistically small fragments. Fusion event is modeled by spatial topology association reactions. There exists certain probability of bond formation when any two nodes are located within two times the node diameter. We designed an iterative simulation scheme to obtain simulations with balanced fission and fusion. First, the system is relaxed and equilibrated for $10^6$ steps without fission/fusion events. Next, initial guesses for the remodeling rates are used to simulate fission/fusion for $2x10^5$ steps. The fission/fusion rates are then updated to compensate for the changes in average fragment size. This cycle is repeated until the changes in average fragment size is small. Lastly, we re-initiate the simulation using the fission/fusion rates established from the latest stable simulation, and adjust the fission/fusion rates every $2x10^5$ steps to compensate any deviation from steady-state dynamics.

To compute the range of MSD accessible to the experimental setup, we used an upper bound of 0.01 $\mu m^2/s$ for node diffusivity $D$ based on the literature[10] and our data, and frame delay $\tau$ ranging from 1 second to 5 second for LLSM. Thus, using $MSD = 6D\tau$, the experimentally relevant MSD is 0.06-0.3 $\mu m^2/s$.

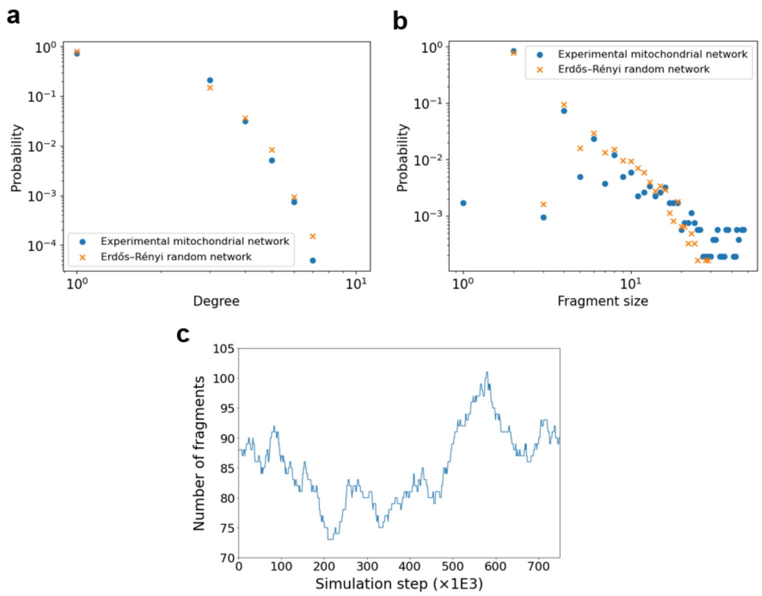


**Supplementary Figure 4. | Network metrics for the mitochondrial simulation. a,b**, The degree distribution (a) and the fragment size distribution (b) for the experimental network segmented by MitoGraph (blue dot), and the random network we generated for simulation (yellow, cross). **c**, The total number of fragments in the simulation is monitored as a function of simulation timestep.

**Supplementary Note 7**

**Fission/fusion detection**

To detect fission and fusion events in the network, we apply a sliding-window approach to identify nodes that undergo persistent structural changes as opposed to transient segmentation variations. First, the fragment indices for each node are recorded for the *half_win_size* frames before and after the current frame, to form the fragment list. Second, for each network edge, the fragment lists for the connected nodes are compared. Fission will be declared if the fragment lists before the current frame are strictly identical, as well as the fragment lists after the current frame are strictly non-overlapping. Since fusion events can be considered as fission events reversed in time, the opposite criterion is used for fusion detection. In the case that multiple remodeling nodes are located in proximity (less than 5 edges away), the nodes are grouped into a single fission/fusion site. At last, the center of the sliding window is moved to the next frame and the computation is iterated.

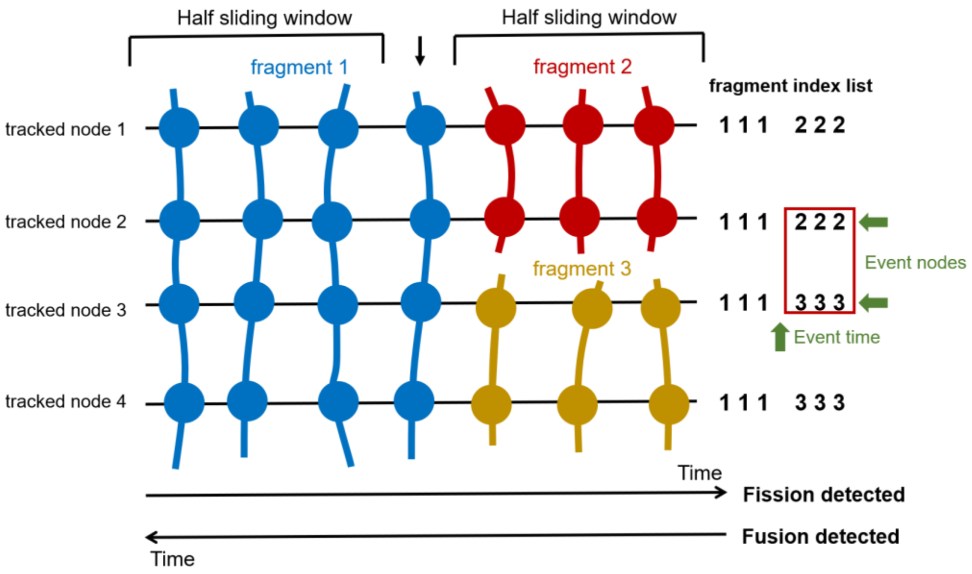


**Supplementary Figure 5. | Algorithm for fission/fusion detection based on node tracking.** Four tracked nodes are positioned vertically, and seven timepoints are shown horizontally. The center of the sliding window is highlighted with the arrow on top. Three fragments are labeled and colored differently. The fragment indices for each node over time are stored. For each half-window, the index values between every two connected nodes are compared frame by frame. An event is declared if fragment indices in one sliding windows are strictly different in time, while those in the other sliding window are strictly identical in time. This requirement is imposed in order to avoid misidentifying transient segmentation noise as remodeling events.

#### Supplementary Note 8

#### Network motility measurements

To measure the network motility of the mitochondrial network, mean-square-displacements (MSDs) vs. time delay curves are computed following previous works[11,12]. The node ensemble-averaged MSDs are plotted for control and oligomycin (Sup. Fig. 6a-b). The MSDs for single tracks are plotted for control and oligomycin (Sup. Fig. 6c-d). Around 80% of the nodes have a coefficient of determination for linear fit greater than 0.8 (Figure 3f). To map diffusivity onto mitochondrial nodes, we first choose the network nodes at a center frame. Next, we collect the node track coordinates 10 frames before and 10 frames after the center frame. In a last step, we compute the MSDs at various time delays in this time window to determine the diffusivity of each node.

To compute vector correlation, we first obtain the displacement vector for a selected node at a selected frame. Next, we compute the vector correlation between this vector and vectors for the nodes directly connected to it, as well as the vectors for the selected node at timepoints 1 frame before and after the selected frame. The vector correlation is given by the dot product of two vectors, divided by the square of the norm of the longer vector. The final correlation value for the selected node at the selected time is reported as the average of these two values.

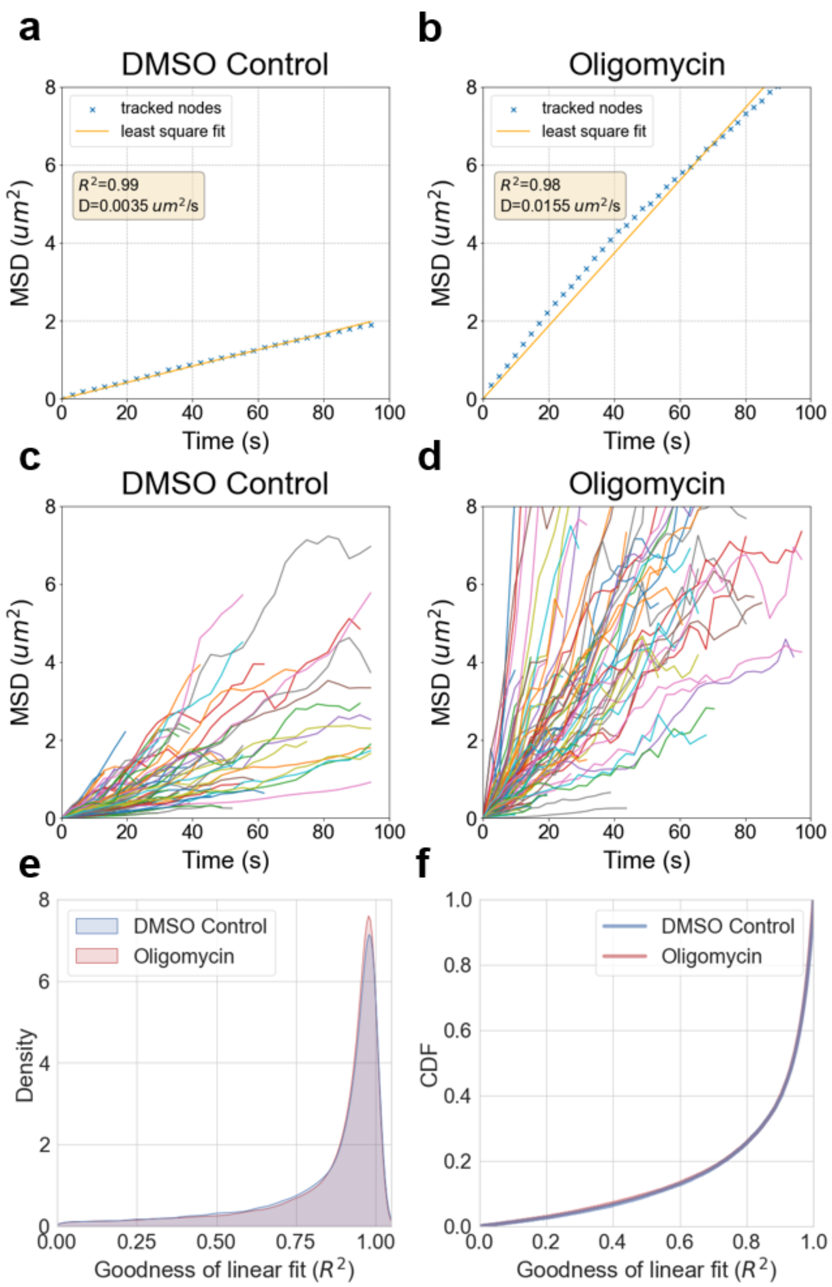


**Supplementary Figure 6. | Mitochondrial skeleton node diffusivity predominantly follows normal diffusive motion. a,b**, Node-averaged mean square displacement (MSD) is computed with respect to time delays (blue) for two conditions, control (a), and oligomycin (b). The linear fit line (orange) is shown together with the coefficient of determination ($R^2$) and diffusion coefficient (D). **c,d,** Node-averaged MSD for DMSO and oligomycin. **e,f**, The goodness of linear fit as measured by $R^2$ is plotted as density distribution (e), and cumulative distribution function (CDF) (f). $R^2$ close to 1 indicates the data points follow a linear pattern.

**Supplementary Note 9**

**Material diffusion simulation on temporal mitochondrial networks**

We use diffusion simulation on temporal dynamic mitochondrial network to understand the flow of material in mitochondrial networks. To calculate global reachability, we initiate a virtual token at every node of the mitochondrial network at the first timestep. Each token is labeled by the source node from which it originated. At every timestep, each node duplicates the tokens it currently has, and transfers them to the connected neighbors. At the end of the simulation, the global reachability for each node is quantified by the number of unique tokens at that node, corresponding to the number of source nodes this node can reach.

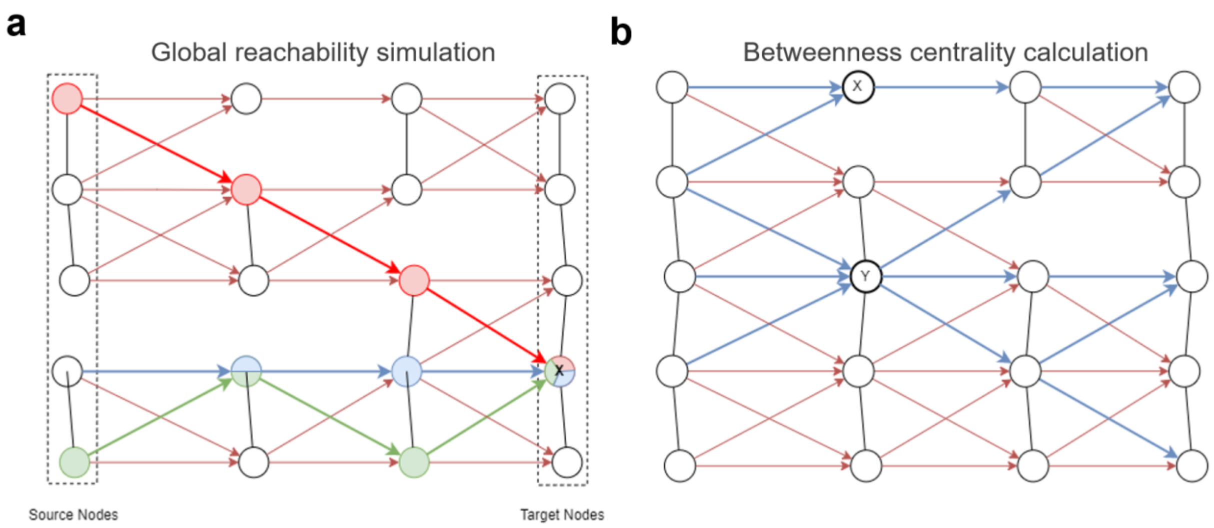


**Supplementary Figure 7. | Schematics for material diffusion simulation on temporal network.** a, Illustration of the global reachability simulation. From each node, material can diffuse into the neighboring nodes or stay in the source node. All possible diffusion pathways over time are marked with an arrow. Time is depicted from left to right. Red, blue, and green arrows indicate representative simulation scenarios. After four timesteps, the target node labeled with X has accumulated three tokens (through the colored transport arrows). b, Illustration of the calculation for betweenness centrality used for determining the central nodes.
